# Improved method for detecting protein-protein interactions using proximity ligation assay

**DOI:** 10.1101/2024.09.01.610697

**Authors:** Zach Tower, Hao Chang

## Abstract

Proximity ligation assay has been widely used to detect protein-protein interaction in cells and tissues. While with great sensitivity, its specificity was often neglected. Here, we report the existence of varying levels of false positives observed with this assay and provide suggestions to minimize false positives for more accurate detection of protein-protein interactions, especially for membrane proteins.

## MAIN

Protein-protein interactions (PPIs) are involved in almost all aspects of biological processes. Studying PPIs is crucial for understanding normal cellular functions, disease mechanisms, and drug discovery. There are several types of methods to detect PPIs in cells. One classic method is the yeast two-hybrid assay (Y2H), which depends on the reconstitution of a functional transcription factor from two domains that are individually fused with two proteins of interest[1]. If two proteins of interest interact, they bring the two domains of the transcription factor together to drive the expression of a reporter gene. The other classic method is co-immunoprecipitation (Co-IP), in which protein complexes are pulled down from cell lysates using an antibody specific to one protein, followed by the detection of the interacting protein using a second antibody[2].

Fluorescence resonance energy transfer (FRET) and bioluminescence resonance energy transfer (BRET) are other types of methods to detect PPIs, which measure the energy transfer between two fluorophores or a bioluminescent donor to a fluorescent protein[3, 4]. When two proteins with fluorescent/bioluminescent tags bind to each other, the energy transfer can be detected by the emission light at a certain wavelength. Proximity ligation assay (PLA) is a recent addition to the toolbox for detecting PPIs[5]. It is based on DNA oligonucleotides-conjugated antibodies that can bind to the proteins of interest. When two proteins of interest bind, the proximity between their corresponding antibodies allows the attached DNA oligonucleotides to be ligated and amplified via a rolling circle mode. The incorporation of fluorescent-labeled oligonucleotides during the amplification generates distinct puncta, which can be analyzed by fluorescence microscopy.

Astrotactin 2 (Astn2) is a transmembrane protein we previously identified as a modifier that regulates mouse hair follicle orientation during development[6, 7]. There are three major membrane proteins that control hair follicle orientations in mice-known as the core planar cell polarity (PCP) proteins. They are encoded by the frizzled family gene *Fzd6* [8], Vangl family gene *Vangl2* [9, 10], and Celsr family gene *Celsr1* [9, 11]. To determine whether Astn2 modulates PCP signaling by directly binding to the core PCP proteins, we applied the widely used Duolink In Situ fluorescent PLA kit to detect their interactions (**Fig. 1a-b**). We transiently expressed the HA-tagged Astn2 with each of the three core PCP proteins (1D4-tagged Fzd6, myc-tagged Vangl2 and Celsr1) in HaCaT cells and performed the PLA following the manufacturer’s instructions. Rabbit anti-HA antibody and mouse anti-1D4 or myc antibody were used to detect the Astn2 and PCP proteins, respectively. Donkey anti-rabbit (+) and donkey anti-mouse (−) probes were used for PLA signal generation. Alexa Fluor 488-conjugated donkey anti-mouse IgG was included to mark the transfected cells. Bright, distinct red fluorescent puncta were seen in transfected cells, but not in non-transfected cells, indicating a strong interaction between Astn2 and all three PCP proteins (**Fig. 1c**).

**Figure 1.**
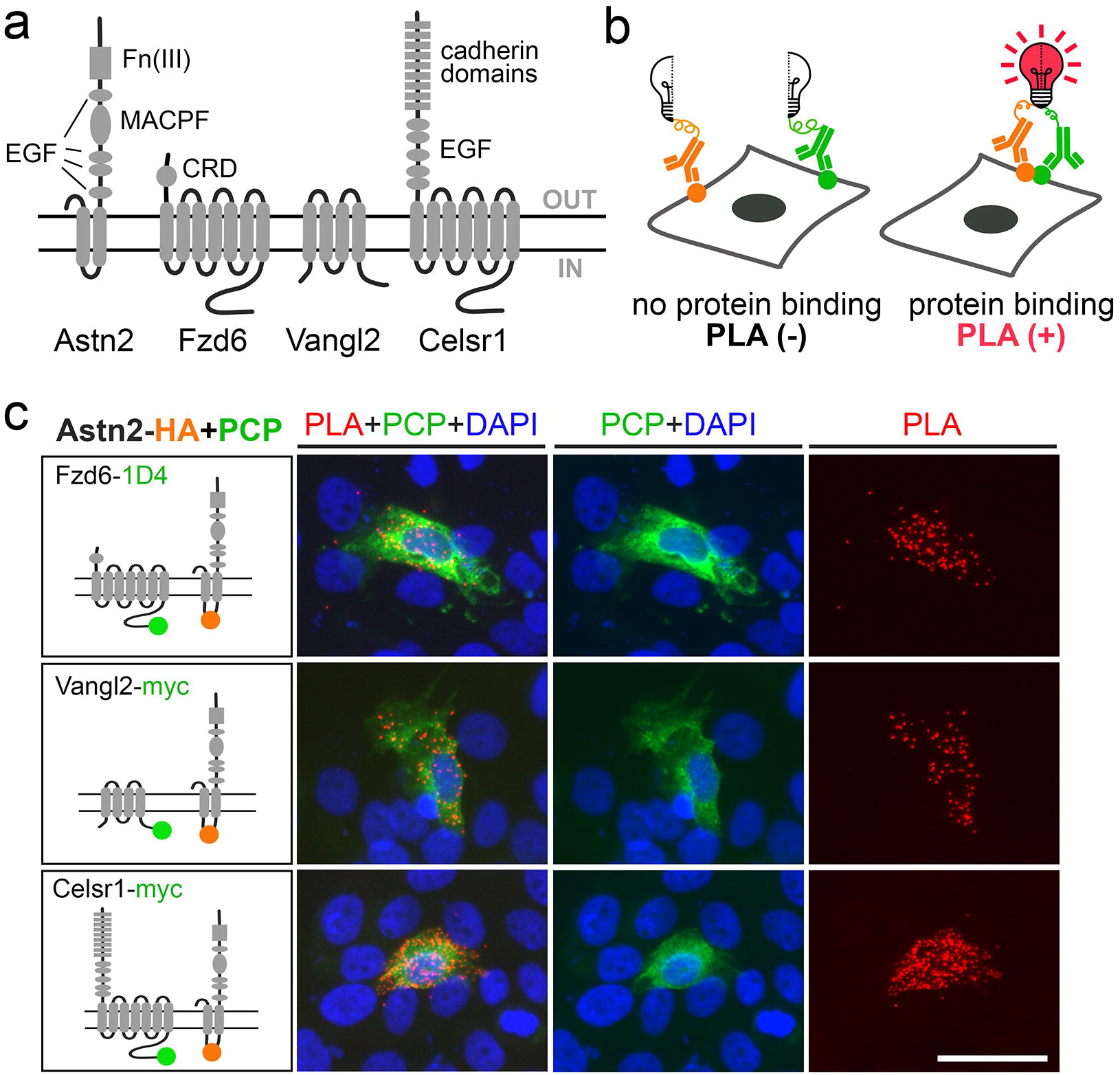
Detection of the protein-protein interactions between Astn2 and PCP proteins using PLA in HaCaT cells. **(a)** Diagram showing the domain structure of Astn2 and three core membrane PCP proteins, Fzd6, Vangl2, and Celsr1. OUT, outside the cell; IN, inside the cell. **(b)** Diagram showing the fluorescent PLA assay. When two proteins of interest bind, a red fluorescent PLA signal will be generated. **(c)** Strong positive PLA signals (red) observed in cells expressing both Astn2 and each of three core PCP proteins (green). Round colored dots in the diagram on the left column mark the position of tags in the tagged proteins. HA in orange, 1D4 and myc in green. Scale bar, 50 μm.

Given that Astn2 has a very long extracellular domain (ECD) at the carboxy-terminus (about 115 kDa, and the entire protein is about 150 kDa), we reasoned that the deletion of the entire carboxy-terminal ectodomain might reduce/abolish the interaction between Astn2 and PCP proteins. We tested this idea with one of the PCP proteins, Fzd6. To generate a potentially more severe perturbation of the interaction, we used the Fzd6 expression plasmid that also lacks the extracellular domain (del ECD). We transiently expressed the HA-tagged truncated Astn2 and 1D4-tagged truncated Fzd6 (with ECD deleted in both plasmids) in HaCaT cells and performed the PLA assay. Strong PLA signals were still observed in cells transfected with these mutant plasmids. We also tested the Fzd6 protein with an intracellular domain (ICD) deletion and found that the deletion of the Fzd6 ICD also does not reduce the PLA signal (**Fig. 2a**). Although we cannot exclude the possibility that the interacting domains reside in regions that we did not delete, e.g., transmembrane domains, it alerted us to test the specificity of the PLA assay with more stringent control experiments.

**Figure 2.**
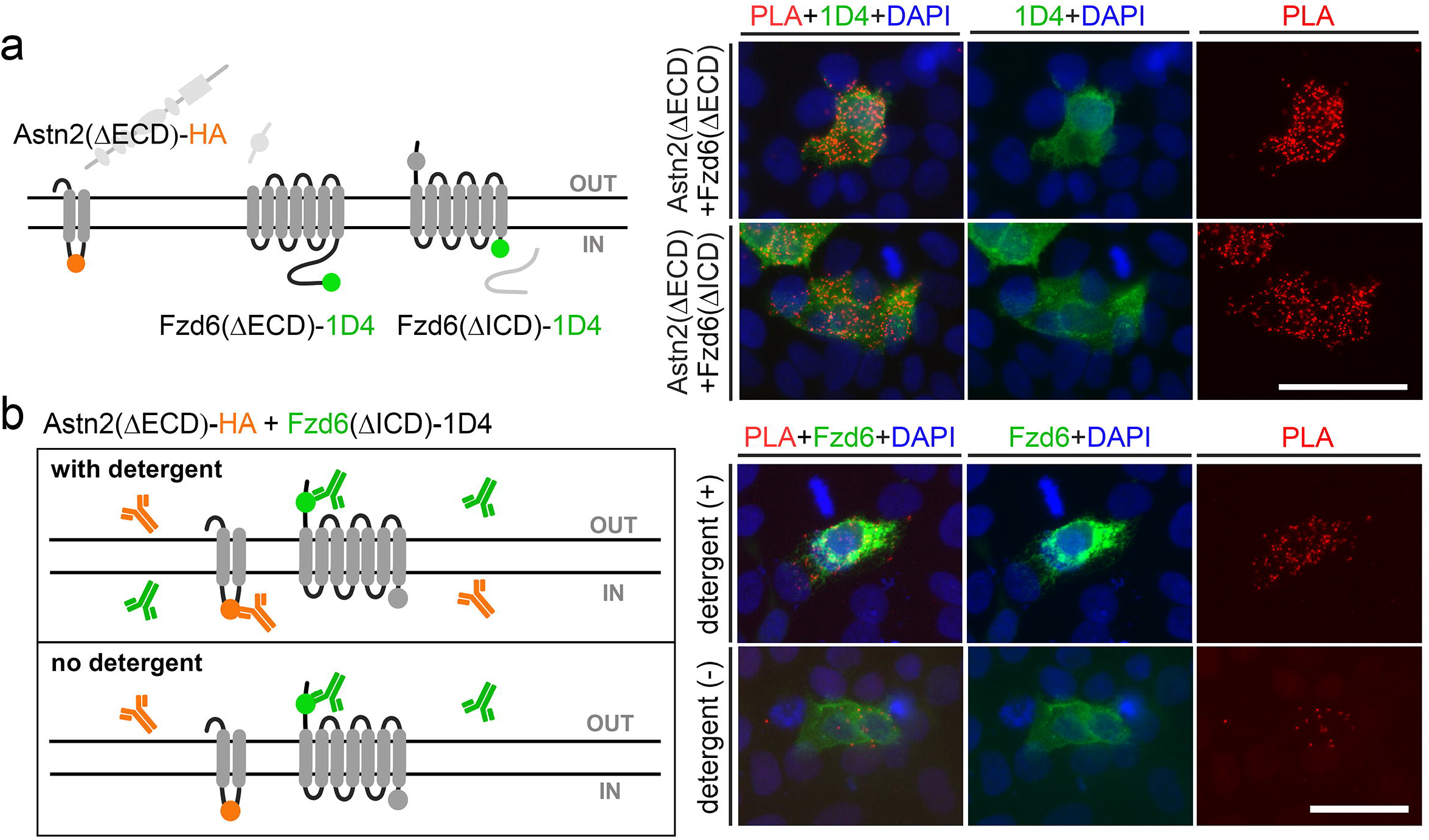
Non-specific PLA signals observed inside and outside the cells. **(a)** Determine the PLA specificity using domain-deleted constructs of Astn2 and Fzd6. Strong PLA signals observed in cells expressing truncated Astn2 (ECD deleted) and truncated Fzd6 (either ECD or ICD deleted). Round colored dots in the diagram on the left column mark the position of tags in the tagged proteins, HA in orange, 1D4 in green. OUT, outside the cell; IN, inside the cell. **(b)** Determine the PLA specificity using two antibodies that bind to different sides of the cell. Strong positive PLA signals (red) were still observed, even without using any detergents during the PLA process. Orange dots and antibodies on the left column indicate the HA tag and corresponding anti-HA antibodies; green dots and antibodies indicate the extracellular epitope inside the endogenous Fzd6 protein and the corresponding anti-Fzd6 antibodies. Scale bar, 50 μm.

We expressed the HA-tagged truncated Astn2 (del ECD) and 1D4-tagged truncated Fzd6 (del ICD) in HaCaT cells and repeated the PLA assay using a different antibody combination. Rabbit anti-HA antibody was used to detect Astn2, as in previous experiments. Instead of using the 1D4 antibody, which recognizes the C-terminal tag of Fzd6 inside the cells, we used a goat-anti-Fzd6 antibody that recognizes the N-terminal extracellular domain of the endogenous Fzd6 protein.

Donkey anti-rabbit (+) and donkey anti-goat (−) probes were used for PLA signal generation. We expected a negative/much reduced PLA signal since the two probes were separated by the cell membrane. The physical barrier of the cell membrane, in addition to the spatial arrangement of the antibodies and the epitopes they bind to, would place the PLA probes outside the proximity range for PLA detection. Surprisingly, we observed an equally strong PLA signal as in the previous experiments when two antibodies used were both localized inside the cells. Next, we repeated the PLA and avoided using any detergents during the staining process. We observed a significant decrease in the numbers of the PLA puncta, suggesting most of the PLA signal in previous experiments with detergents came from inside the cell, not on the cell membrane where the protein interaction should naturally occur. Without detergents, only one antibody (Fzd6) can bind to the cell surface. The presence of strong PLA puncta, although in reduced numbers, suggests a certain background level of noise with this assay (**Fig. 2b**). We should point out that we never observed any distinct PLA puncta in cells transfected singly with either Astn2 or Fzd6 expression plasmid, suggesting the non-specific PLA signal we observed is not relevant to the antibody usage, staining protocol, or detection method.

In conclusion, we demonstrated that strong PLA signals can be robustly generated when applying the antibody-based fluorescent PLA assay to study PPIs of membrane proteins. A clear contrast of positive and negative results (e.g., hundreds of puncta in double transfected versus zero in single or non-transfected cells) can also be observed in the same experimental settings. However, caution must be taken, given our observation that non-specific false positive signals can occur in double-transfected cells. Based on our findings, we suggest: (1) Cells should always be co-stained at least for one of the proteins during the PLA to ensure they indeed express the proteins of interest. PLA staining kits with triple fluorescent channel options are commercially available (green/red channels for co-staining two individual proteins and far-red channel for PLA detection). This will eliminate the false negative result caused by the no/low expression of target proteins. (2) Mutant proteins with domain deletions should be included as the negative control, whenever possible. This will eliminate the false positive of PLA and ensure two proteins indeed bind to each other. (3) When studying PPIs of membrane proteins, use antibodies that recognize endogenous epitopes located outside the cell or tag the proteins of interest with an extracellular tag. Avoid using detergents when performing PLA – this will significantly reduce the non-specific signals that occur inside the cells. This background reduction strategy obviously cannot be applied to intracellular proteins since antibodies need detergent treatment to penetrate the cells.

## METHODS

### Plasmids and cell culture

Expression plasmids for wild-type and truncated Astn2 and Fzd6 were described previously[6, 12]. Vangl2 and Celsr1 expression plasmids were kindly provided by Dr. Jeremy Nathans at Johns Hopkins University. HaCaT cells were kindly provided by Dr. Jun Dai at the University of Wisconsin-Madison. Standard procedures were used to transfect plasmid DNA into HaCaT cells grown in DMEM medium with 10% fetal bovine serum. For each well of cells to be transfected in a 12-well plate, 0.5 μg of DNA was diluted in 50 μl of serum-free media. 1.8 µl of 1 mg/ml polyethylenimine (PEI) was added into the diluted DNA solution, mixed gently, and incubated for 10 minutes at room temperature before adding to the cells. Cells were incubated for 48 hours post-transfection before assaying.

### PLA assay

HaCaT cells were fixed with formalin at 4°C overnight and washed with PBS. For the regular PLA assay (with detergents), we strictly followed the instructions of Duolink PLA fluorescence protocol (Sigma). Briefly, cells were washed in PBST (0.3% Triton X-100) for 10 minutes to permeabilize the membrane, blocked with Duolink Blocking Solution for 1 hour at 37 °C, and incubated with primary antibodies overnight at 4 °C. The following primary antibodies were used: rabbit anti-HA (#3724, Cell Signaling Technology, 1:1000), mouse anti-1D4 (#MA1-722, ThermoFisher, 1:1000), mouse anti-myc (#2276, Cell Signaling Technology, 1:1000), and goat anti-Fzd6 (AF1526; R&D Systems; 1:400). Duolink rabbit (+), mouse (−), and goat (−) probes were used for ligation and amplification. Alexa Fluor 488-conjugated donkey anti-mouse or goat IgG antibodies (Invitrogen, 1:1000) were also included in the probe incubation step to co-stain the PCP proteins. Duolink In Situ Mounting Media with DAPI was used to mount the cells.

Images were captured using a Biotek Lionheart fluorescent microscope with Gen5 software. For the PLA assay without detergents, the following adjustments were applied. Cells did not undergo PBST wash before the PLA assay. 5% normal donkey serum in PBS was used for blocking and diluting the primary antibodies instead of the Duolink Blocking Solution and Antibody Diluent.

## ACKNOWLEDGMENTS

The work was supported by the National Institutes of Health grant R01GM129259 (to HC). HC was also supported in part by the Cripps/Ratcliff Professor for Skin and Cancer Research Endowed Professorship and the Department of Veterans Affairs VA Merit Review Award I01CX002308.

## AUTHORS’ CONTRIBUTIONS

Conceptualization: HC; Methodology: ZT, HC; Formal analysis: ZT, HC; Investigation: ZT, HC; Data Curation: ZT; Writing: ZT, HC; Project administration: HC; Funding acquisition: HC.

